# A hypercubic Mk model framework for capturing reversibility in disease, cancer, and evolutionary accumulation modelling

**DOI:** 10.1101/2024.06.27.600959

**Authors:** Iain G. Johnston, Ramon Diaz-Uriarte

## Abstract

Accumulation models, where a system progressively acquires binary features over time, are common in the study of cancer progression, evolutionary biology, and other fields. Many approaches have been developed to infer the accumulation pathways by which features (for example, mutations) are acquired over time. However, most of these approaches do not support reversibility: the loss of a feature once it has been acquired (for example, the clearing of a mutation from a tumour or population). Here, we demonstrate how the well-established Mk model from evolutionary biology, embedded on a hypercubic transition graph, can be used to infer the dynamics of accumulation processes, including the possibility of reversible transitions, from data which may be uncertain and cross-sectional, longitudinal, or phylogenetically / phylogenomically embedded. Positive and negative interactions between arbitrary sets of features (not limited to pairwise interactions) are supported. We demonstrate this approach with synthetic datasets and real data on bacterial drug resistance and cancer progression. While this implementation is limited in the number of features that can be considered, we discuss how this limitation may be relaxed to deal with larger systems.

## Introduction

Many systems of interest in biology, medicine, and beyond involve the progressive, random, coupled accumulation of different binary features over time. Such features – variously called traits or characters – may be, for example, the presence or absence of particular mutations in a tumour (Diaz-Uriarte and Johnston 2024; Schill et al. 2020; Beerenwinkel et al. 2015; Schwartz and Schäffer 2017), susceptibility or resistance to particular drugs in evolving pathogens (Greenbury, Barahona, and Johnston 2020; Moen and Johnston 2023; Beerenwinkel et al. 2005), the presence or absence of genes in evolving genomes (Johnston and Williams 2016), or the presentation or absence of particular clinical symptoms in patients (Johnston et al. 2019; Schill et al. 2024). In all these cases, the study of the dynamics by which features are acquired – the trajectories or pathways of accumulation – are of interest in both describing the scientific system and predicting its future behaviour.

The field of accumulation modelling considers how sets of discrete, usually binary, features are acquired over time in a system. Many approaches have been developed in the cancer literature, where features are often driver mutations, and samples are independent patients (Schill et al. 2020; Diaz-Uriarte and Herrera-Nieto 2022; Beerenwinkel et al. 2015; Montazeri et al. 2016; Diaz-Uriarte and Johnston 2024; Angaroni et al. 2022; Schill et al. 2024) or, recently, trees reflecting the development of tumours over time (Luo, Kuipers, and Beerenwinkel 2023; Aga et al. 2024; Caravagna et al. 2018). There are 2^L^ possible patterns of presence or absence for each of L features, and transitions are usually considered to correspond to the acquisition of exactly one feature. Here, a system may be in one of 2^L^ states, and transitions between any pair i and j of these states occur with a rate r_ij_, which can be nonzero only if j differs from i by the presence of one feature.

The Markov k-state (Mk) model (Pagel 1994; Lewis 2001) is widely used in evolutionary biology to infer the dynamics by which features (usually called characters, in this context) change between discrete states on a phylogeny. Here, every instance of a system can occupy one of k discrete states, with Markovian transitions allowed between these states. As an evolutionary process branches into different lineages, descendants inherit their ancestor’s state and may transition to other states independently of their sister lineages. Different versions of the model place, or relax, different constraints on the rates of the transitions between states, which are parameters of the model. The original real-world example is how two binary traits, multiple mating partners and posterior decoration, co-evolve in primates (Pagel 1994). More generally in the Mk model (Boyko and Beaulieu 2021), a system may be in one of k states, and transitions between any pair i and j of these states occur with a rate r_ij_ for a given instance of the system.

As discussed previously (Moen and Johnston 2023), a mathematical connection exists between these two approaches, clearly visible from the final sentences of the preceding two paragraphs. If k = 2^L^ and the transition rates for the Mk model are constrained to be zero for the necessary pairs of states (those that differ by the acquisition of more than one change), the Mk model becomes the accumulation model (Fig. 1, A.1 and A.3).

**Figure 1.**
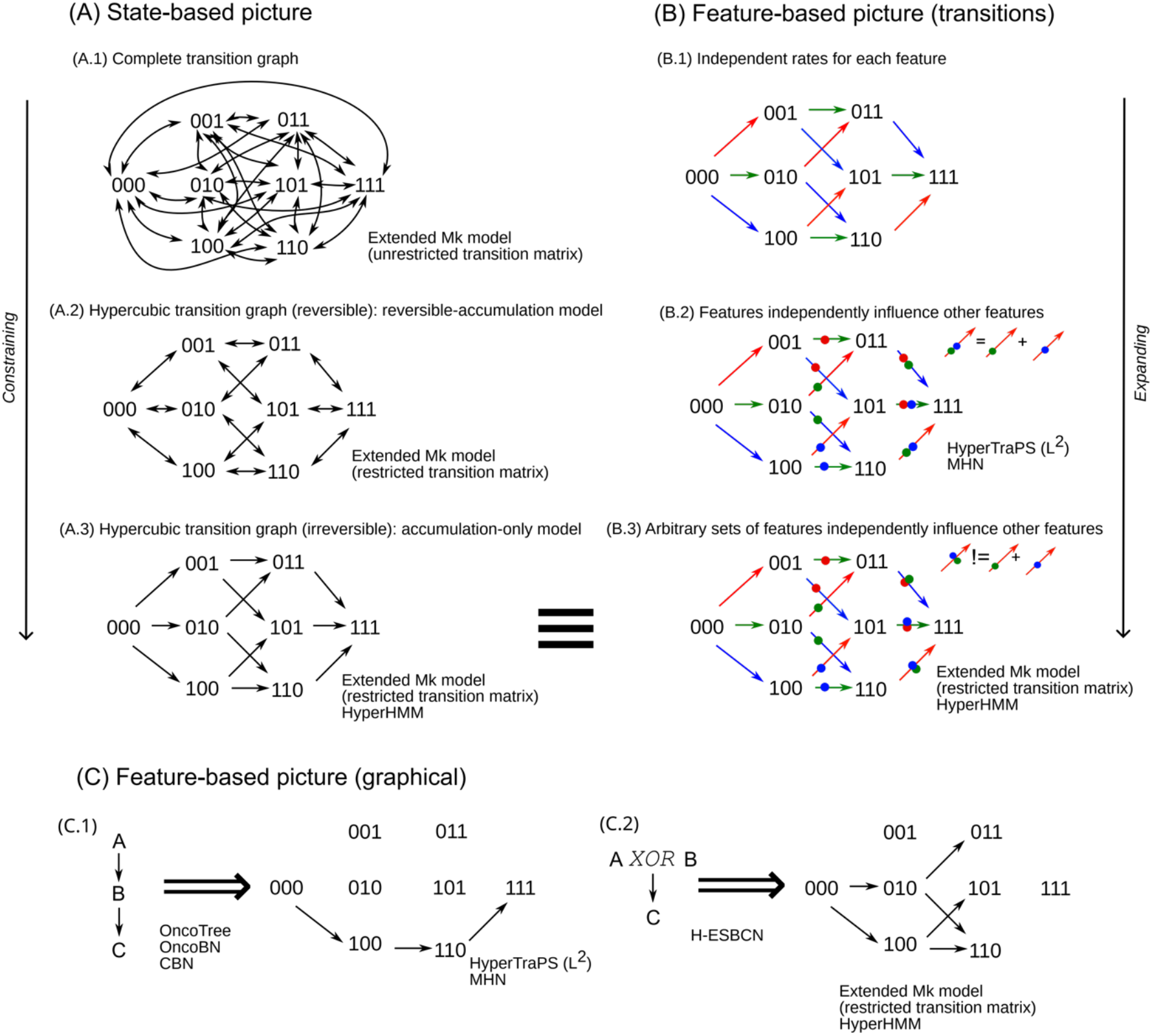
State spaces and features in (reversible) accumulation and Mk models. (A) Transitions supported in increasingly constrained modifications of the Mk model, with rates based on *states:* A.1 unrestricted dynamics, A.2 reversible accumulation, A.3 irreversible accumulation. (B) Increasingly flexible modifications of the accumulation model with rates based on *features and their interactions*. Model B.1 (L^1^ rates) has the same rate (colour) for every transition involving a given feature; model B.2 (L^2^ / mutual hazards) allows pairwise influences between features, so that having acquired feature X can influence the rate of acquiring feature Y (denoted by a coloured ball for each acquired feature); model B.3 allows arbitrary contributions to all feature rates from any subset of acquired features. Models A.3 and B.3 are equivalent. (C) Correspondence between the Directed Acyclic Graph (DAG) representation of feature-based models and their hypercubic representation. In C.1, the DAG describing the dependence of feature acquisitions is linear: C depends on B, which depends on A. In C.2, C depends A and B via a logical XOR relationship. This panel also shows two common alternative notations: letters to denote the altered (mutated) feature in DAGs vs. 0/1 to indicate non-altered/altered for the corresponding feature in hypercubic graphs.

Consider now a *reversible* accumulation model, which is the same as the accumulation model, but where r_ij_ is now also allowed to be nonzero if j differs from i by the *absence* of one feature. In other words, the system can transition by acquiring *or losing* individual features. This reversibility challenges existing approaches designed to model accumulation processes, because an infinite set of accumulation pathways now in principle exists between any two states (involving arbitrarily long sequences of feature gain-loss pairs). However, this reversible picture still corresponds to a subset of the more general Mk model (Fig. 1, A.1 and A.2). Indeed, using a (potentially hidden) Markov modelling framework to explore the coupled evolution of many binary traits is a central idea behind evolutionary analysis packages like *corHMM* (Boyko and Beaulieu 2021). Here we will show that existing methods from the evolutionary literature for inferring parameters in the Mk model can readily be applied to this reversible accumulation picture, allowing analysis of this problem.

The value and necessity of including reversibility in accumulation modelling of course depends on the scientific question. In some cases, the mechanisms of the underlying process mean that the loss of features can be safely neglected. Whether this is the case depends both on the scientific field and the entity being considered in the modelling process. We will explore these points further in the discussion, and proceed by introducing an approach by which reversibility can be included – and tested against an irreversible picture – as and when scientifically desired.

## Methods

### State spaces in (reversible) accumulation and Mk models

Consider a state space **S** = {S_0_, …, S_2_^L^_-1_ }, consisting of 2^L^ states. For convenience, we set the index of a state to the decimal value of a binary string of length L, corresponding to the absence (0) or presence (1) of each of the L features in that state (for example, for L = 3, 000 ≡ 0, 001 ≡ 1, 010 ≡ 2, …, 110 ≡ 6, 111 ≡ 7). This is a general procedure for coding combinations of binary states as multistate characters that has been used before with the Mk model (for example, in (Boyko and Beaulieu 2021), p. 470). This transformation allows us to seamlessly use the Mk model for accumulation processes that include the cross-sectional case, and that, then, allow us to model both irreversible and reversible cases.

We label the transition rate from the state with index i to the state with index j as r_ij_. A general accumulation-only model allows r_ij_ > 0 if and only if j-i = 2^k^, where k is a non-negative integer. In other words, transitions are only supported from i to j if i lacks exactly one feature that j possesses. The corresponding transition graph for this model takes the form of an L-dimensional hypercube with single directed edges (Fig. 1, panel A.3). A reversible accumulation model allows r_ij_ > 0 if and only if j-i = 2^k^ or i-j = 2^k^. Hence, two (different) nonzero rates can correspond to the transition from i to j (gaining a feature) and from j to i (losing that feature). Now the transition graph is an L-dimensional hypercube with pairs of opposing directed edges between vertices (Fig. 1, panel A.2).

Now consider an Mk model with k = 2^L^ states. The most general picture (called the extended Mk model) allows r_ij_ > 0 for all i and j. Hence, the transition graph is a complete graph, with pairs of opposing directed edges between every pair of vertices (Fig. 1, panel A.1). Clearly, both the transition graphs for the accumulation-only and reversible accumulation models are subgraphs of the extended Mk model transition graph, and can be captured by constraining the required set of edges in the Mk model to be zero.

### Relationships between observations in (reversible) accumulation and Mk models

We now consider the forms of data that can be used in the different modelling cases. Accumulation modelling often considers cross-sectional, independently observed samples (motivated by the cancer literature, where data has traditionally consisted of single observations for independent patients). Some developments including HyperTraPS (Greenbury, Barahona, and Johnston 2020; Johnston and Williams 2016; Johnston and Røyrvik 2020) and recent expansion HyperTraPS-CT (Aga et al. 2024), HyperHMM (Moen and Johnston 2023), and TreeMHN (Luo, Kuipers, and Beerenwinkel 2023) allow observations to be connected via a tree. This may exactly be a phylogenetic tree in evolutionary applications, or a phylogenomic tree (corresponding to different cell lineages evolving within a patient or within a tumour) for disease cases. In all cases, the typical assumption is that a precursor or ancestral state exists which has not acquired any of the features considered (the patient before any cancer developed, for example).

The Mk model considers phylogenetically embedded data, where observations are often taken to represent the tips (terminal vertices) on a phylogenetic tree. This situation is motivated by the evolutionary biology literature, where we often observe properties of modern-day samples that are historically connected by their evolutionary relationship. The state of the root of the tree, and of internal nodes, is not in general known precisely (being in the unobserved evolutionary past) and must be inferred if required.

These various cases can be collected under an overall umbrella (Fig. 2). A dataset is connected by a set of n_tree_ trees. In the cross-sectional case, we can either picture n_tree_ = 1 “star” trees connecting a single root node with state 0 to n independent observations, or n_tree_ = n “stump” trees, each connecting a root at state 0 to a single descendant observation; for cross-sectional data, the single star tree, with possibly different branch lengths is appropriate where different branch lengths reflect different times to sample. For the other cases, the trees (phylogenetic or phylogenomic) connecting observations come directly from the data.

**Figure 2.**
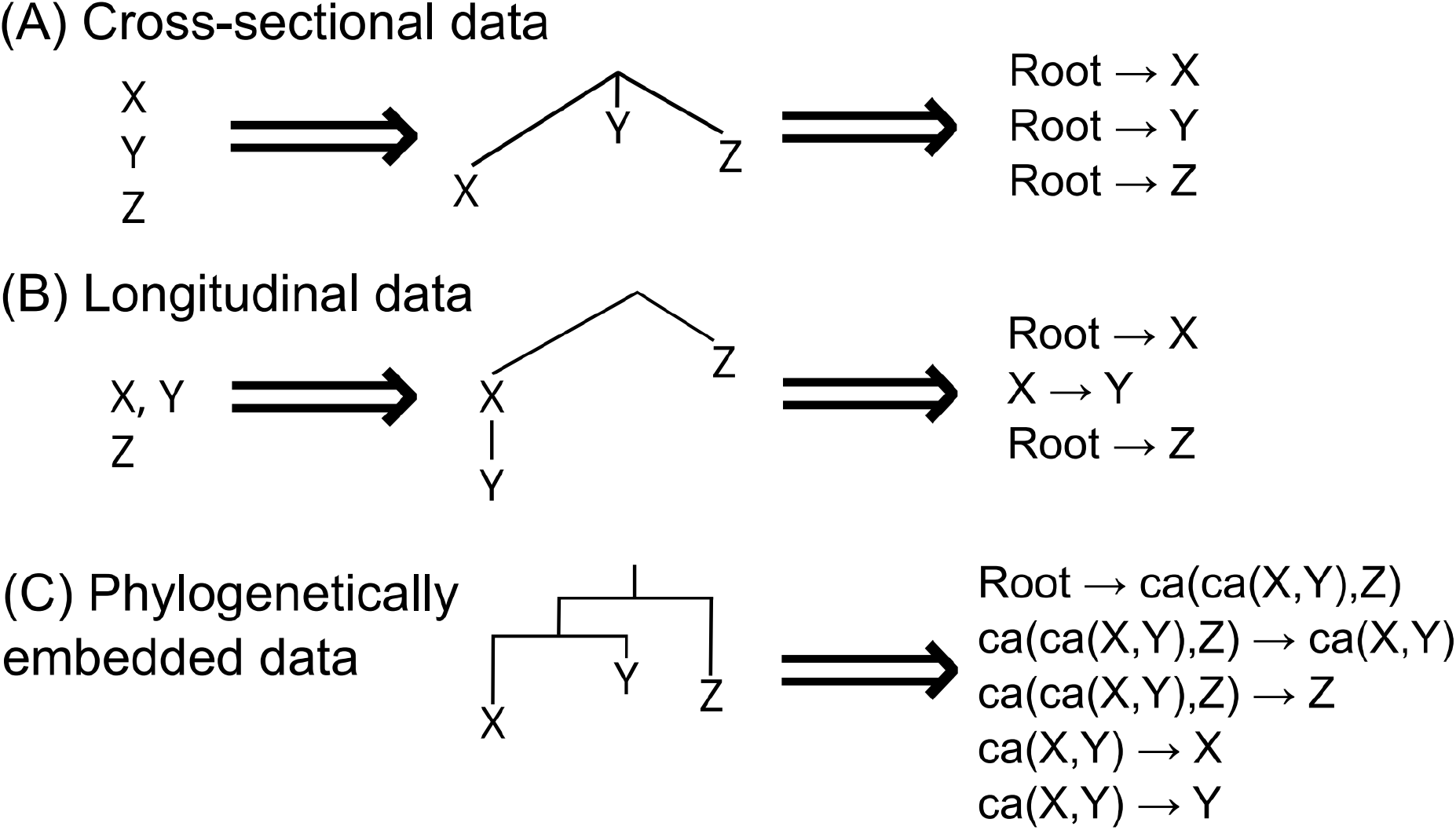
Data types in accumulation modelling, and casting them as trees. (A) Cross-sectional data. Independent observations X, Y, Z can be represented as independent lineages branching from an initial state (the root of a polychotomous tree). The source data are then the set of independent transitions from the root state to each observation. (B) Longitudinal data. Each independent series can be represented as an independent lineage branching from an initial state, with progressive observations in a series following that lineage. The source data are then transitions from the root to each lineage’s ancestor, then transitions within the lineage. (C) Phylogenetic data. With a phylogeny, transitions are independent ancestor-descendant pairs. Ancestral states (ca, common ancestor) can be reconstructed using phylogenetic methods and/or invoking assumptions about the dynamics of the particular evolutionary process.

We will refer to this picture, using a reversible or irreversible hypercubic transition graph in conjunction with the Mk model, as the “hypercubic Mk model” or “HyperMk” for short, but emphasise that this refers to the model structure and not a particular numerical implementation. As we discuss below, different implementations are possible and further refinement for the HyperMk case specifically are certainly feasible.

### Numerical implementation

Given data, inference of the transition parameters in an Mk model can be performed via likelihood maximization, often using Felstenstein’s pruning algorithm (Felsenstein 1973), a dynamic programming approach recursing up the levels of the tree and considering conditional dependencies on ancestral states. Many specific software implementations exist for this process (see below and Supp. Fig. 1). We will focus on the *castor* package in R (Louca and Doebeli 2018; Louca and Pennell 2020) as an easily-implementable, open-source, solution with the flexibility to deal with the different HyperMk structures described above. The Mk fitting routine in *castor* (function `fit_mk`) supports inference using multiple trees and constraints on transition parameters, as well as the specification of prior states for unobserved vertices.

Data that is already embedded on one or more trees can naturally be passed to *castor*. Cross-sectional data could, in principle, be passed as a set of “stump” trees with one root and one tip. However, we found that this protocol challenged the numerical behaviour of the fitting procedure. Instead we adopt the following protocol. For every observation, we construct a tree with one root and two tips. Using the capacity of *castor* to support prior distributions on vertices, we enforce that one tip must have the state corresponding to observation i, and allow the second tip to be completely unconstrained (uniform prior over all states). In this way we produce a branching tree structure that is readily analysed by *castor* without introducing any artefacts into the dataset.

Several other R packages offer the ability to fit Mk models of a given structure. The *corHMM* package (Boyko and Beaulieu 2021) offers a powerful interface for inferring transition parameters of (hidden) Markov models, which can readily be used to implement this hypercubic Mk model picture. The representation of states as binary strings is an explicit focus of that research, making the hypercubic modelling framework particularly easy to implement. However, we could not find a straightforward way of analysing cross-sectional or multiple independent trees using this package. *phytools* (Revell 2012) also includes functionality for fitting Mk models and allows the natural inclusion of cross-sectional observations by considering a “star” phylogeny (polychotomous independent branches from an initial state). Here, we are not using *phytools*’ fitPagel function to model correlated evolution because that function only allows for two traits, and because our transformation of combinations of binary traits into multistate characters is the standard procedure for analyzing with multiple, possibly correlated characters (Methods, (Boyko and Beaulieu 2021)). However, the inclusion of uncertain data, and the adjustment of the parameters of the underlying optimization process, were less straightforward in this approach than in *castor*; we also found that some observations were apparently ignored in some cases. We underline that these – and likely more – alternatives may readily be used for many of the case studies in this report; Supp. Fig. 1 demonstrates their application to a simple case study, and the code repository includes examples of using these different libraries as the core solver.

The code implementing this setup in R (R Core Team and Team 2022) is freely available at https://github.com/StochasticBiology/hypermk and in addition to the libraries described above also uses libraries *ape* (Paradis and Schliep 2019) and *phangorn* (Schliep 2011) for modelling trees, *ggplot2* (Wickham 2016), *ggraph* (Pedersen 2020), *igraph* (Csardi and Nepusz 2006), *ggtree* (Yu et al. 2017), and *ggpubr* (Kassambara 2020) for visualization.

### Pruning and model comparison

In both the irreversible and reversible models, it can (and often is) the case that many edges in the hypercubic transition network reflect transitions which do not contribute to the likelihood. For example, in the irreversible case, if 000-001 is the only transition from the initial state 000, no transitions from 100 or 010 will ever contribute to the likelihood, as those source states are unreachable from the initial state. We implemented a “pruning” method to remove parameters associated with these negligible transitions (Supp. Fig. 2). Pruning – removing particular edges from the hypercubic transition graph -- can readily be used to enforce a particular subset of dynamics in a model (Supp. Fig. 2B), but can also be used to simplify models by removing these extraneous parameters (Supp. Fig. 2C). From a fitted model, we simulate dynamics on the transition network and track the probability associated with each transition. We then explicitly remove edges with probability below a threshold (by default 0.01, reduced to 10^−5^ for the tuberculosis case study to avoid removing edges that contribute to rare observations), and refit the model without these parameters. For all cases we confirm that the “pruned” likelihood is not detectably different from the original likelihood, confirming that we have not removed edges that contribute to the likelihood. The “pruned” version of that model, with a likelihood and parameter count, can be used in comparisons via model selection criteria; we use AICc, corrected AIC, by default (Supp. Fig. 2).

## Results

### Synthetic case studies

To test the capacity of the Mk model picture to capture accumulation models, we construct a set of illustrative case studies with synthetic data. We first simulate a phylogeny using a birth-death process and then simulate an evolutionary process on this phylogeny, involving L=5 features; the different instances discussed below differ in the accumulation model. In the first, irreversible, instance, these features are acquired in the deterministic order 00000-10000-11000-11100-11110-11111. The corresponding tree is shown in Fig. 3A. The Mk model, with parameters restricted to support irreversible accumulation, immediately identifies the generating pathway (Fig. 3B-C). For the illustrated dataset, the AICc of the HyperMk model with irreversible dynamics is 123.8, and that with reversible dynamics is 129.4; the lower AICc for the irreversible model reflects the fact that the less complex irreversible model is sufficient to capture these accumulation-only dynamics.

**Figure 3.**
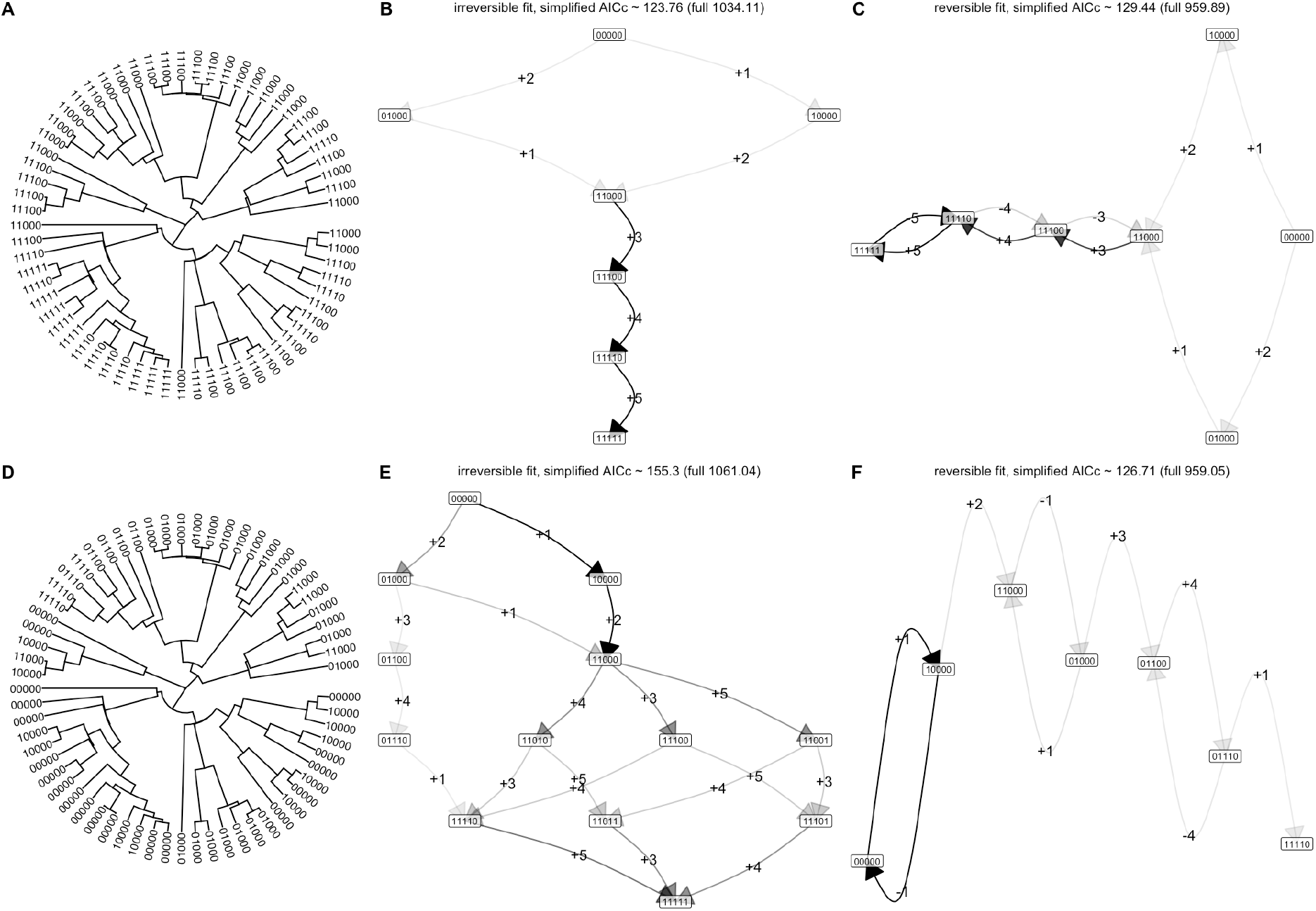
Capturing irreversible and reversible accumulation dynamics with the Mk model. (A) Observations of different states of an irreversible process on the tips of a 64-node randomly-simulated tree. (B) Inferred transition graph for these data, using an irreversible model. Node labels give different states; edge labels give the feature that is gained (+) or lost (-) at each transition, but in this irreversible case no feature losses (-) are allowed. The darkness of an edge gives its rate: darker edges have higher associated transition rates. (C) Inferred transition graph for these data using a reversible model. The AICc values for the full fitted model, and for the simplified model discarding all edges that do not support flux from the 00000 state (i.e., edges with two or more transitions), are given: the AICc for the irreversible model is lower, reflecting the irreversible generating process. (D) Observations of different states of a reversible process on the tips of a 64-node randomly-simulated tree. Here, the first feature can be lost as well as gained. (E) irreversible and (F) reversible model transition graphs as in (B-C). In the reversible case, the “-1” edges corresponding to the loss of the first feature appear ubiquitously; the AICc value for the reversible model is now lower, reflecting the model’s better ability to capture reversible dynamics.

Next, we allow the first feature to undergo reversion, so that transitions involving the loss of the first feature are allowed from any state (Fig. 3D). The corresponding outcomes are shown in Fig. 3E-F. Now, the reversible model has a lower AICc (126.7 compared to 155.3), reflecting the fact that a model allowing reversible transitions better capture the observations than the accumulation-only model.

To demonstrate HyperMk’s ability to capture positive and negative influences between traits, and to work with cross-sectional as well as phylogenetically-embedded observations, we next consider an L=4 case where two pathways are supported: 0000-0001-0011-0111-1111 and 0000-1000-1100-1110-1111. These two pathways are mutually repressing, so that the first step taken from 0000 completely determines the subsequent dynamics. We use the protocol described in Methods to represent observations from this system as a collection of trees that can be analysed by *castor*, and show the results in Fig. 4A-C. As expected, given the lack of restrictions placed on the transitions, HyperMk readily captures the cross-repressing nature of these pathways.

**Figure 4.**
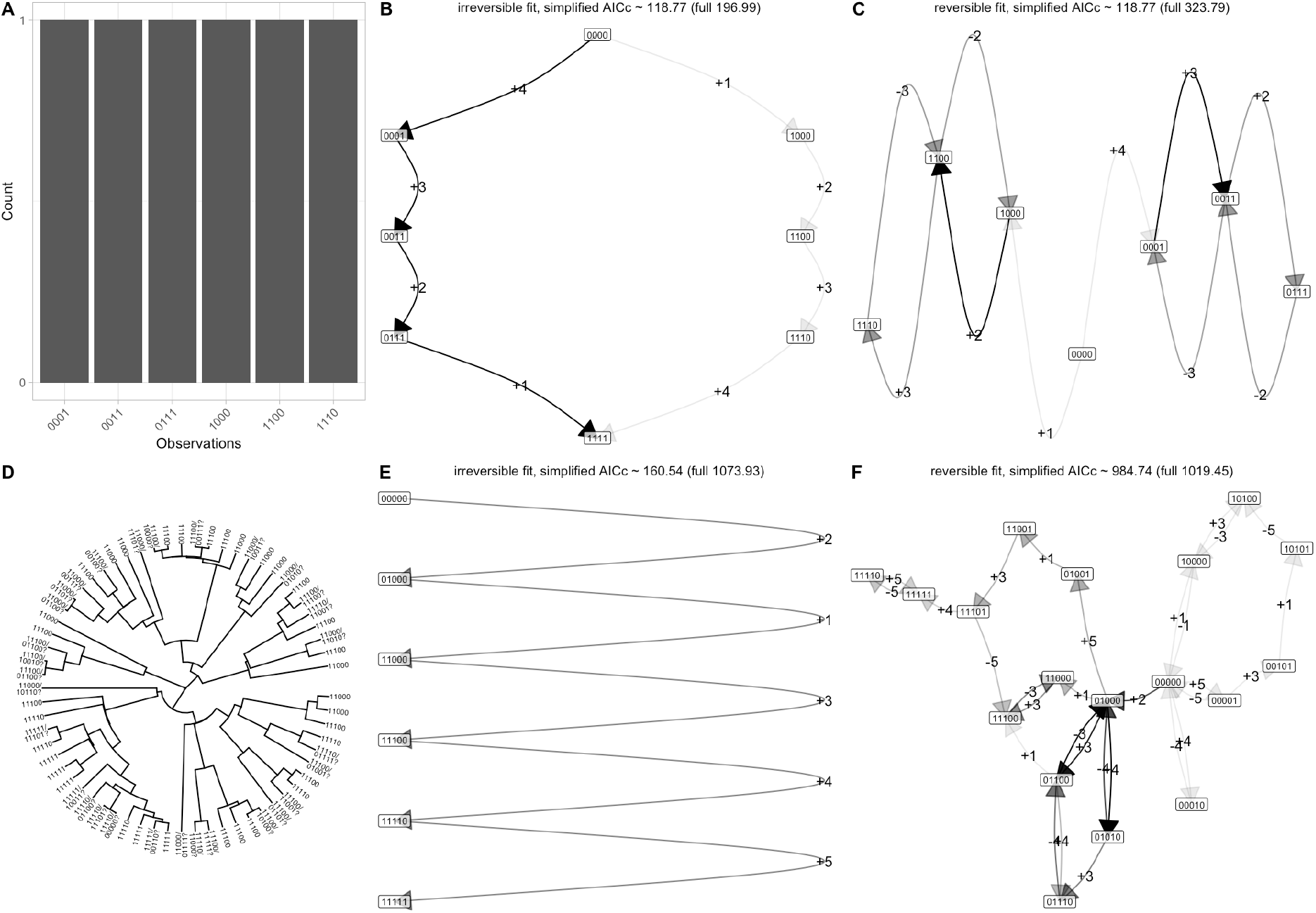
Capturing positive and negative influences, cross-sectional observations, and uncertainty with the Mk model. (A) Cross-sectional observations generated by the two-pathway system described in the text. Now no tree links observations, which are taken to reflect independent samples. (B-C) Inference of transitions graphs for this cross-sectional case using (B) irreversible and (C) reversible models. In both cases, the competing pathways (features 4, 3, 2, 1 versus features 1, 2, 3, 4) are readily distinguished. (D) Observations on a simulated tree, where some observations are uncertain. Here, the single pathway (features 1, 2, 3, 4, 5) generates observations, but half of the observations (marked “?”) are confounded by mixing with another random state. (E-F) Inference of transition graphs for this uncertain case using (E) irreversible and (F) reversible models. The core pathway is recovered in both models but with some probability flux to other states supported.

The Mk model can naturally account for uncertainty in observations, and uncertainty in the inferred transition parameters can be quantified via resampling, for example through using the bootstrap. In the fit routine in the *castor* package, uncertain observations are included by specifying “tip priors”: P_is_ is the likelihood of the observed state of tip i given that it was truly in state s. We demonstrate inference using data generated from the single pathway model (Fig. 3A-C) with uncertainty in Fig. 4D-F. For the irreversible model, HyperMk readily identifies a parsimonious generating mechanism; for the reversible model, the core of this mechanism is identified but with corresponding uncertainty in the form of some probability of transitions involving states that are not generated by this process^1^.

### Cancer progression

Although reversibility of genetic changes in cancer progression is likely less important than in other systems, we also include a case study of cancer progression to connect with previous studies. We use an old but commonly-used dataset on chromosomal aberrations accumulating during ovarian cancer progression (Knutsen et al. 2005). These samples are taken from different patients and we therefore use the cross-sectional implementation of HyperMk. We focus on L=5 features, labelled by chromosome, chromosomal arm, and gain or loss of genetic material: 8q+ means a gain of material q arm of chromosome 8. In the order presented in the data, the feature set is (8q+, 3q+, 5q-, 4q-, 8p-) (Fig. 5A shows the frequency distribution of the chromosomal aberration combinations).

**Figure 5.**
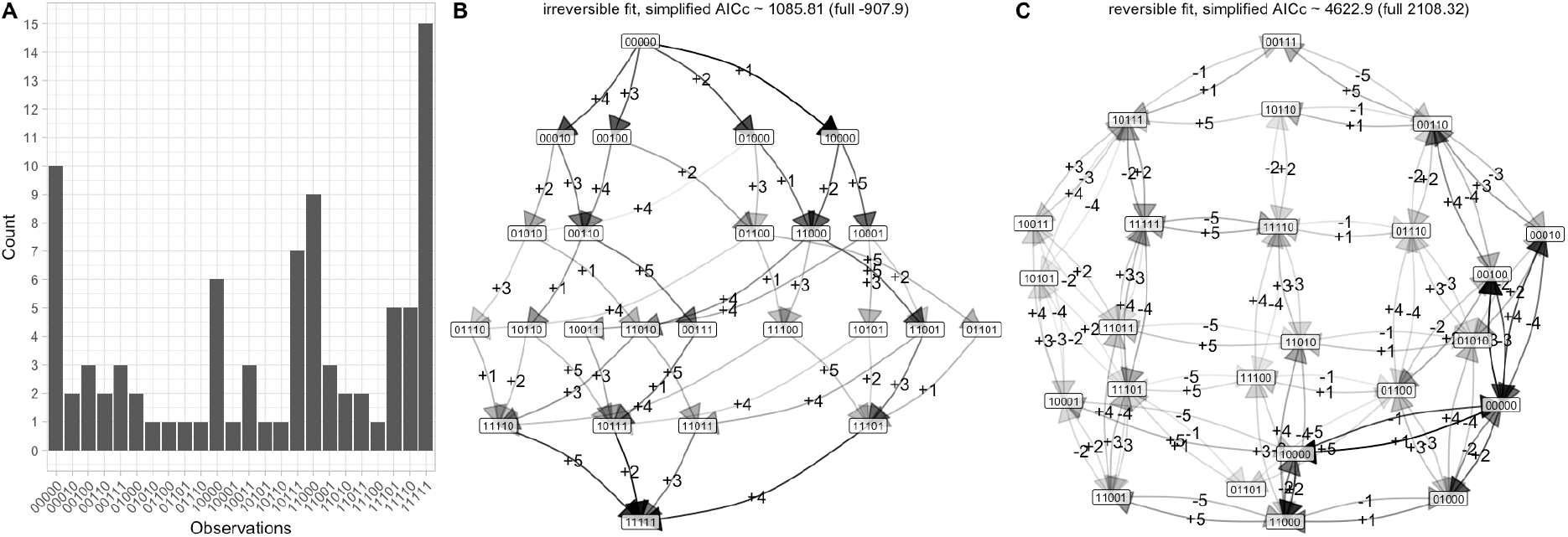
Accumulation dynamics in ovarian cancer progression. (A) The set of 87 cross-sectional observations of chromosomal aberrations in ovarian cancer (from independent patients, not linked by a phylogeny) from (Knutsen et al. 2005). The (reduced) set of aberration locations is (8q+, 3q+, 5q-, 4q-, 8p-). (B-C) Inference of transition graphs using (B) irreversible and (C) reversible models.

The most common pathways consistently identified by previous analyses have 8q+ and 3q+ as first steps, with 5q- and 8p- as competing early next steps and 4q-occurring on average somewhat later. Average orderings from HyperHMM (Moen and Johnston 2023) place 8q+ with the highest probability first (lower probability 5q- and 3q+), 3q+ second (lower probability 8p- and 5q-), 8p-, 4q-, or 8q+ third, 5q-, 4q-, or 8q+ fourth, and 8p-fifth (lower probability 5q-). These patterns are all borne out by both the reversible and irreversible inferred transition graphs (Fig. 5B-C), which support several pathways of progression dominated with 8q+ and 3q+ as first steps, with the other transitions occurring in consistent orderings thererafter.

Agreeing with the intuition that reversible transitions may play a limited role in determining cancer progression pathways (see Discussion), the AICc of the irreversible model is lower than that of the reversible model, suggesting that accumulation-only dynamics is the preferred picture for this dataset. The loss transitions supported by the inferred reversible model largely mirror acquisition transitions with lower probability (for example, a lower-probability -1 transition from 10000 to 00000 mirroring the +1 transition from 00000 to 10000), suggesting that unmatched loss transitions do not play a substantial role in determining dynamics (unlike, for example, the loss of feature 1 from multiple different states in the synthetic reversible example above, Fig. 3D-F).

### Anti-microbial resistance

To demonstrate the application of HyperMk to real-world questions, we consider data on the evolution of anti-microbial resistance in *Mycobacterium tuberculosis* (TB) (Casali et al. 2014), as curated in (Greenbury, Barahona, and Johnston 2020). Here, we take 150 isolates of TB with associated profiles of drug resistance, describing susceptibility (absence) or resistance (presence) to a collection of drugs. The observations are phylogenetically connected via a tree that was characterized using genetic data in the original publication (Fig. 6A). For this illustrative case we consider the first L=5 of these drugs: isoniazid (INH), rifampicin/rifampin (RIF), streptomycin (STR), ethambutol (EMB), pyrazinamide (PZA). Previous work using accumulation-only models HyperTraPS and HyperHMM (Greenbury, Barahona, and Johnston 2020; Moen and Johnston 2023) have shown that of this set, INH and RIF are the most likely first acquisitions, followed by EMB then STR.

**Figure 6.**
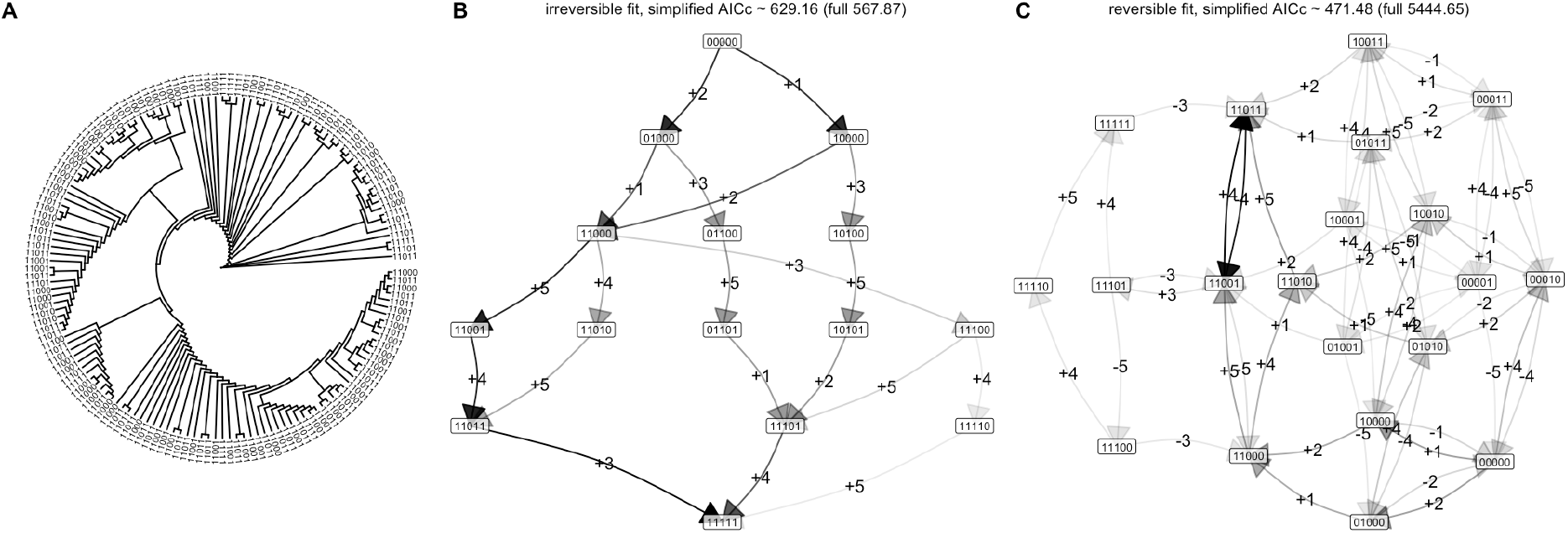
Evolutionary dynamics in tuberculosis drug resistance. (A) Observations of patterns of drug resistance from (Casali et al. 2014). The phylogeny is plotted with uniform branch lengths for legibility; the true phylogeny has heterogeneous branch lengths. The (reduced) set of drug features is (INH, RIF, STR, EMB, PZA); 0 corresponds to susceptibility, 1 corresponds to resistance. (B-C) Inference of transition graphs using (B) irreversible and (C) reversible models.

Following HyperMk fitting, this ordering is clearly visible in both reversible and irreversible models (Fig. 6B-C). The reversible model experiences a clear advantage according to AICc (471.5 vs. 629.2 for the reversible and irreversible models, respectively), with the transition graph showing substantial probability flux associated with the loss of STR and EMB resistances.

### Implementation

The runtime of this approach, implemented using *castor*, does not scale particularly tractably with the number of features L. Fig. 7 shows runtime in seconds for a fit involving N observations each of L features, on a simulated tree as in Fig. 3. Every additional feature multiplies the runtime by at least one order of magnitude, and sometimes approaching two. Feature sets larger than L=7 will take inconveniently long to infer with this implementation. However, as discussed below, this implementation both uses a “full” parameterization where no reduction of parameter space is used, and a general Mk model fitting package. Indeed, one key contribution of this paper is showing the connection between the different problems represented in Fig. 1, something that is evidenced by using off-the-shelf Mk software for all of them. A custom implementation for specific problems, involving, for example, only pairwise interactions between features and/or specific speedups from the hypercubic transition network structure, may afford substantial gains in speed. As described in Methods, several different existing packages in addition to *castor* provide functionality for fitting Mk models compatible with this accumulation modelling picture (Supp. Fig. 1).

**Figure 7.**
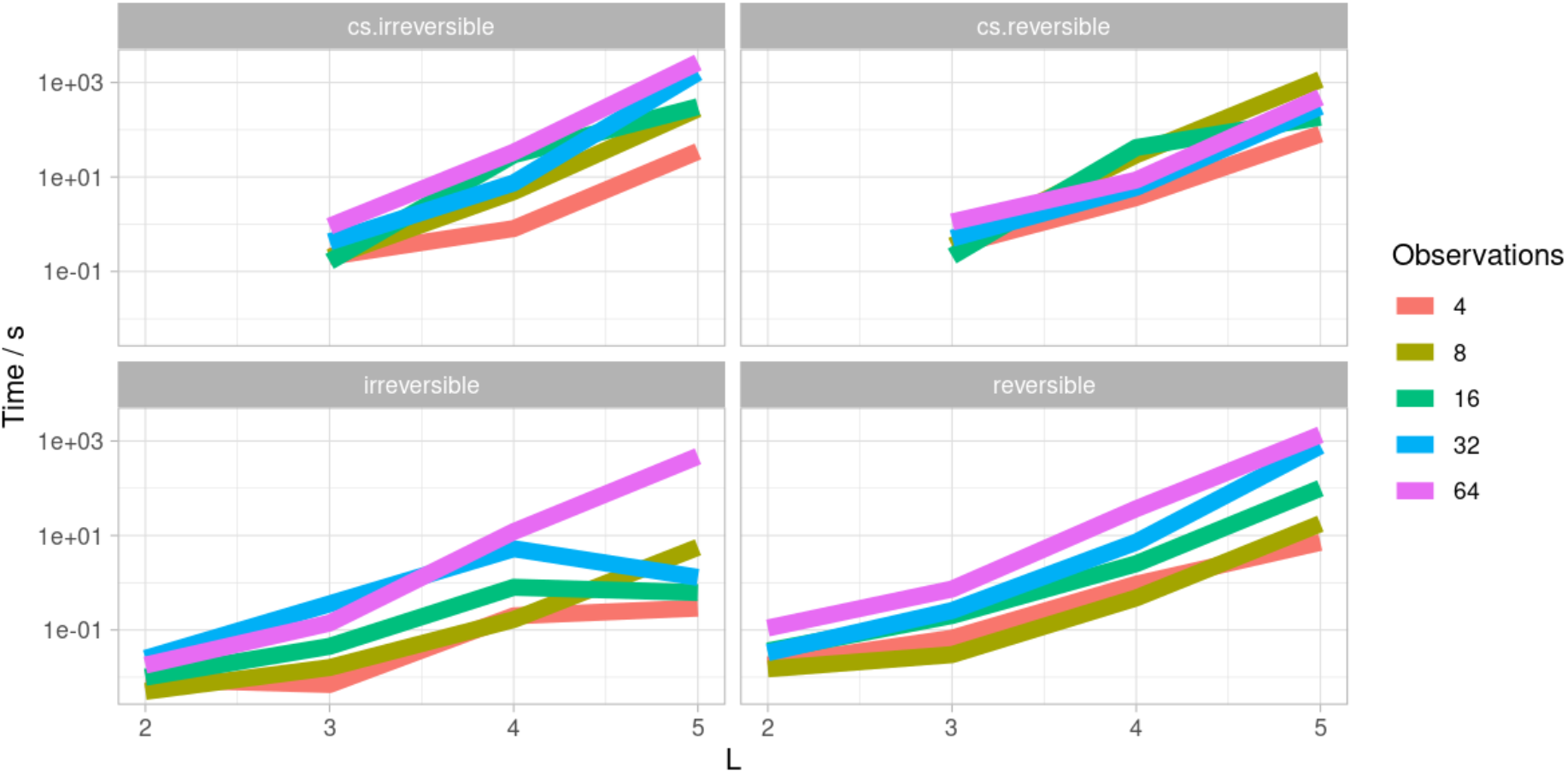
Timing for synthetic case studies. Simulated fits using *castor* to implement HyperMk. Data are randomly generated, either from a uniform samples of states (cross-sectional examples) or from a simulated tree and a reversible generating process as in Fig. 3A. Timings are for a single 3.2 Ghz core (Apple M1 Pro).

We observed that numerical convergence is a consistent issue with these approaches. Intuitively, the presence of reversible transitions means that an infinite set of possible pathways can in principle be responsible for a given observation.

While in practice any non-infinite transition rates will act to decrease the probability of longer pathways, this dramatic expansion of possibilities over the irreversible case makes inference with the reversible model numerically challenging. We found that a combination of multiple start points and looser constraints on the number of permitted iterations mitigated these challenges to some extent – at the inevitable cost of more computational resource. There were also some library-specific issues related to this convergence: the Mk model fit with *castor* sometimes crashed when multiple start points were invoked. Our working approach – implemented in the software accompanying this article – is thus to avoid multiple start points and only parallelise over different experiments; the case studies here in Figs. 3-6 are all tractable on a single machine, with the tuberculosis case study taking the longest (several hours) to converge.

## Discussion

We have introduced and demonstrated a way of supporting reversible transitions in accumulation modelling using the Mk model. Of course, the value of supporting reversibility depends on the scientific question being asked. In cancer progression, mutations occur and clonally proliferate as cells divide. Depending on what the “entity” being modelled is – a cell, a tumour, a patient, etc (Diaz-Uriarte and Johnston 2024) – reversibility may simply be inappropriate to consider. The probability of a reverse mutation “undoing” a particular mutation at the cellular level is presumably very low, although at the tumour level it is not inconceivable that cell-level selection could remove a mutation from the cellular population once it has arisen.

In evolution, for example the evolution of anti-microbial resistance, this question can be case-specific. AMR traits in *Klebsiella pneumoniae* are often acquired via transfer of plasmids – mobile DNA elements – which can be gained and very readily lost again (Holt et al. 2015). AMR traits in tuberculosis are often acquired via chromosomal mutations, which are less readily lost once they have been acquired (Casali et al. 2014). At the level of single cells, reversibility may therefore be more important to consider for Klebsiella than tuberculosis. And – in agreement with Fig. 6 – reversible dynamics may occur at the population level (one strain outcompeting another to be the dominant observed type) without requiring single-cell reversibility (Diaz-Uriarte and Johnston 2024). In other cases, reversibility may be simply ill-posed – for example, in accumulation modelling of students completing tasks in online courses (Peach et al. 2021).

We have not focussed on a continuous time variable in this study, instead viewing dynamics as “ordinal” and reporting the sequence, rather than timing, of transitions. This approach mirrors HyperTraPS and HyperHMM (Greenbury, Barahona, and Johnston 2020; Moen and Johnston 2023; Johnston and Williams 2016), but many other approaches (including a recent generalization of HyperTraPS (Aga et al. 2024) assume a continuous timescale that is implicitly or explicitly connected to the observations (Schill et al. 2020; Diaz-Uriarte and Johnston 2024; Schill et al. 2024; Diaz-Uriarte and Herrera-Nieto 2022). The Mk model approach described here has the full capacity to capture continuous timing in observations, as the branch lengths in the phylogeny (or in the set of synthetic trees, for cross-sectional data) correspond to times between states. The rate parameters learned in the model fitting then become true rates in the time behaviour of the system, rather than just describing relative propensities of events. Accumulation problems where continuous time data are available can exploit this natural feature of the Mk approach.

A clear shortcoming in this implementation is the number of features that can be considered. Existing methods for accumulation modelling can deal with dozens, and sometimes over a hundred, interacting features – although the case of arbitrary dependencies between sets of features, not just pairwise interactions, is more limited (see details and discussion in Aga et al., 2024). Here, feature sets larger than L=7 approach computational intractability. However, it is very possible that different software implementations will change this limitation – and future developments exploiting the restrictions on parameter space specific to the reversible accumulation model may substantially increase efficiency. In particular, the picture outlined in this research allows arbitrary independent transitions between states, rather than coarse-graining transition parameters by the features involved in the transition (Fig. 1). More restricted parameterisations, for example reflecting the pairwise-only interactions used in the L^2^ parameterisation of HyperTraPS and equivalent in Mutual Hazard Networks, would lead to smaller parameter spaces and presumably faster inference. The *corHMM* package (Boyko and Beaulieu 2021) allows different classes of parameter structure to be specified; implementations of accumulation modelling for single trees (or a more general picture, with future development) may provide a straightforward solution for this adaptation.

More generally, connections between the evolutionary and systematic literatures and accumulation modelling may allow more transfer of promising insights between the fields. One scenario that is likely to become more prevalent with single-cell sequencing involves data sets where several patients provide multiple within-patient samples which are phylogenetically related (Luo, Kuipers, and Beerenwinkel 2023; Aga et al. 2024; Caravagna et al. 2018) which naturally fit in the hypercubic Mk modelling framework. A broader common question in applied accumulation modelling is whether continuous and/or ordinal features, as well as presence/absence features, can be analysed in the same framework. While binary accumulation approaches have been used to analyse datasets containing ordinal features (for example, the levels of coma presentation in malaria patients (Johnston et al. 2019)), this has typically been done with a rather crude coding approach involving inequalities (a set of n features denoting “presence” of x < 1, x < 2, x < 3, and so on). A recent advance in the systematics literature provides an approach by which the coupled dynamics of discrete and continuous characters can be naturally analysed (Boyko, O’Meara, and Beaulieu 2023). Broadly, we anticipate that if the usual picture of data linked by a dichotomous phylogenetic tree with branch lengths can be relaxed – to star phylogenies or sets of trees or “stumps” for example – a wide range of approaches from phylogenetic methods will find use in the analysis of accumulation processes.

## Acknowledgements

This project has received funding from the European Research Council (ERC) under the European Union’s Horizon 2020 research and innovation programme (Grant agreement No. 805046 (EvoConBiO) to IGJ). IGJ was supported by the Trond Mohn Foundation [project HyperEvol under grant agreement No. TMS2021TMT09], through the Centre for Antimicrobial Resistance in Western Norway (CAMRIA) [TMS2020TMT11]. RDU was supported by grant PID2019-111256RB-I00 funded by MCIN/AEI/10.13039/501100011033.

## Supplementary Figures

**Supplementary Figure 1.**
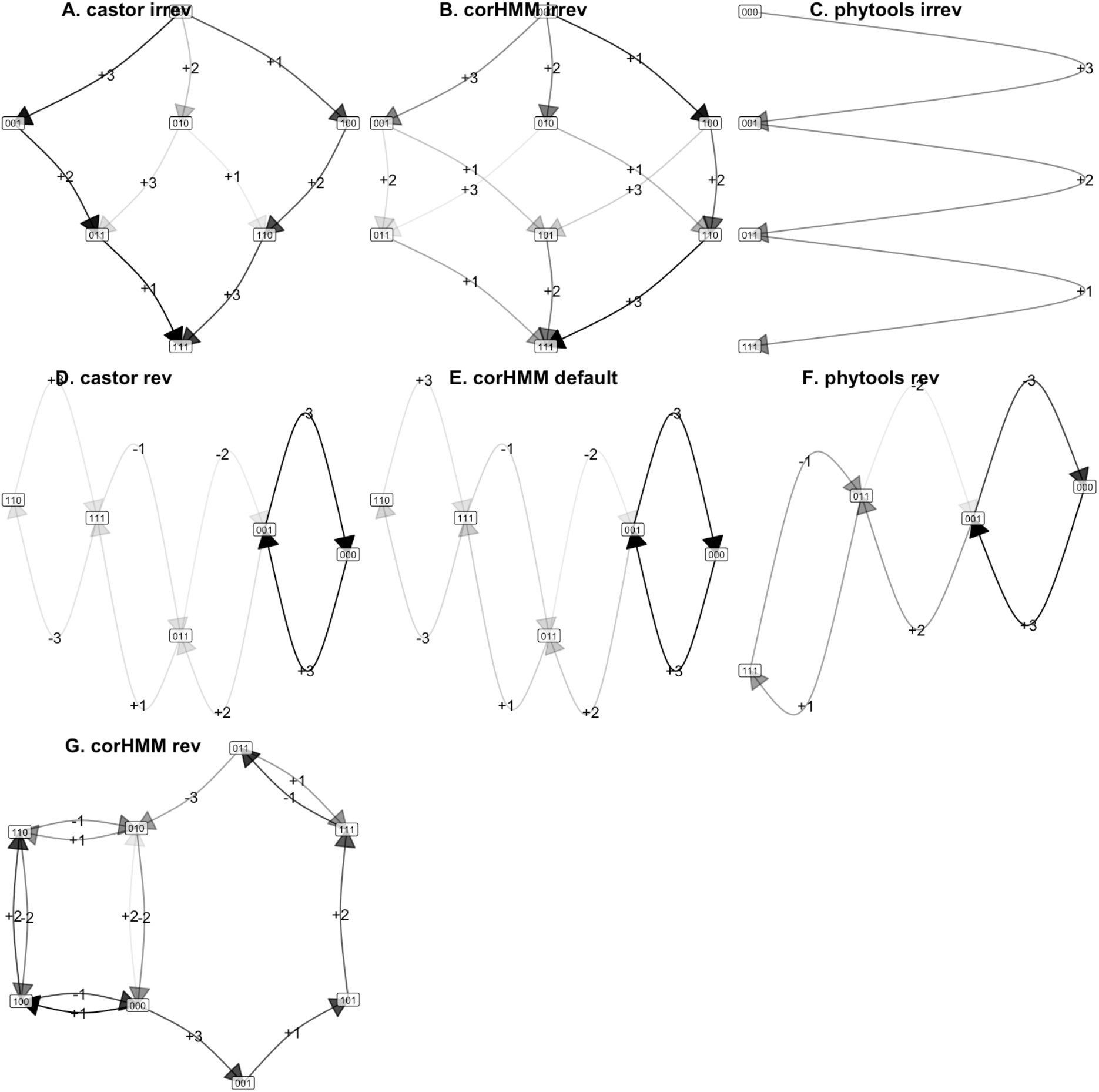
Different Mk model fitting approaches. Inferred transition networks for synthetic data capturing a single pathway with reversible dynamics (observations 000, 001, 011, 111, 110). (A-C) irreversible model fits using the single pathway case study using *castor, corHMM*, and *phytools*; (D-F) reversible model fits. The *phytools* fit in this implementation seems to ignore the 110 observation. In (E) the default *corHMM* fit has been performed, which considers only states present in the observation set; in (B, G) the presence of all 2^L^ possible binary states has been enforced.

**Supplementary Figure 2.**
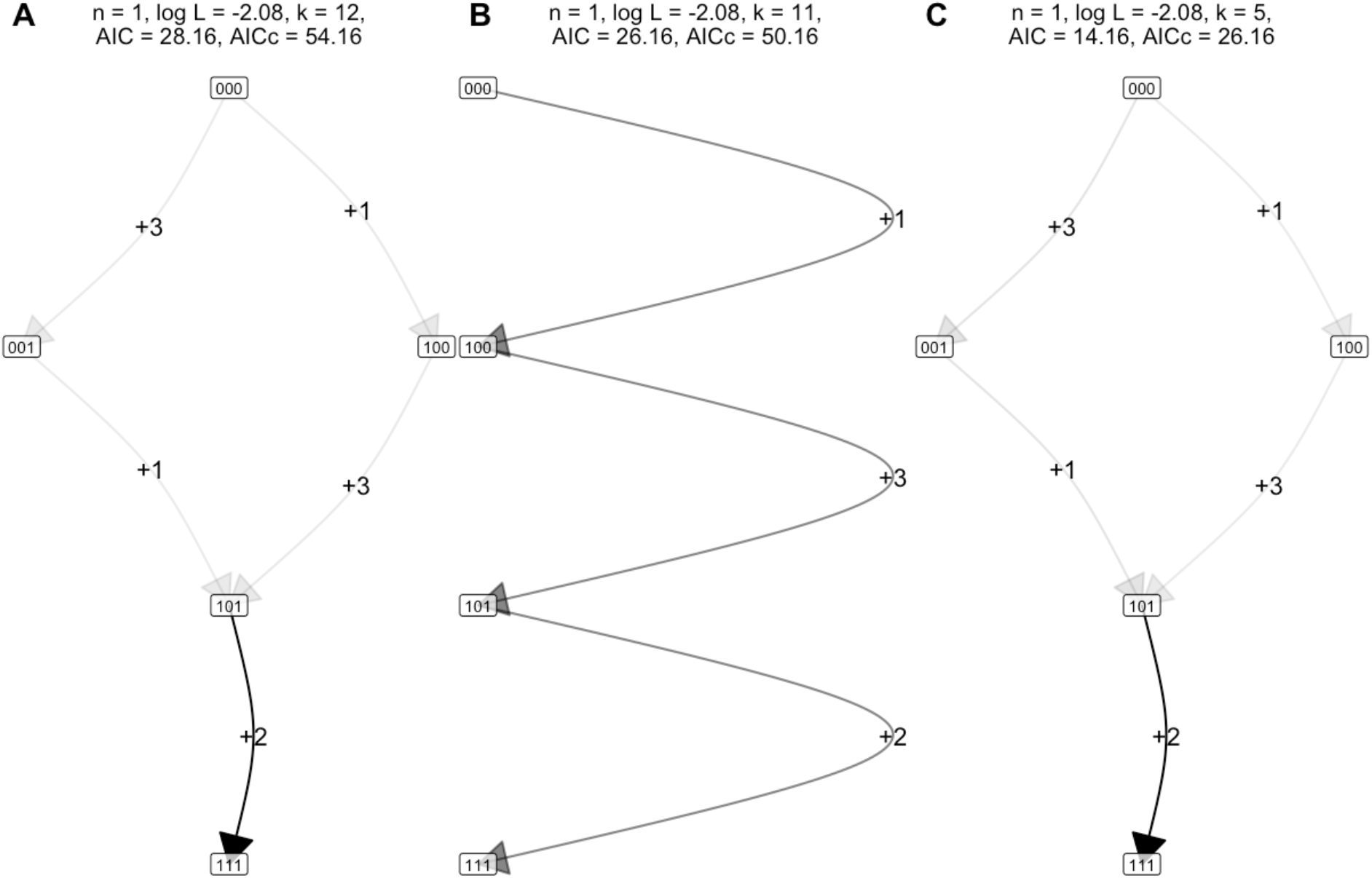
Illustration of “pruning” parameters. (A) An unpruned fit of an irreversible L = 3 model to a single (n = 1) observation 101 involves a model with k = L 2^L-1^ = 12 edge parameters. Only those 5 supporting nonzero flux in the fitted model are plotted here; the others are not identifiable from the data. Equal weight is given to the possibilities 000-001 and 000-100, both of which are compatible with the observation. (B) The edge parameter 000-001 has been removed and the model refitted; now the dynamics are completely constrained and the AIC is lowered by 2 (2× number of parameters removed). (C) All parameters corresponding to fluxes < 10^−4^ in (A) have been removed and the model refitted. Differences to the inferred dynamics are negligible and the AIC is lowered by 14 (2× number of parameters removed) compared to (A). This method (C) is used throughout the study.

## Footnotes

1 The *castor* package also supports bootstrap resampling and multiple, parallelised fitting attempts per bootstrap, though we have not made use of it here.

## Notes

https://github.com/StochasticBiology/hypermk

### Competing Interest Statement

The authors have declared no competing interest.

## References

Aga, Olav NL, Morten Brun, Konstantinos Giannakis, Kazeem A. Dauda, Ramon Diaz-Uriarte, and Iain Johnston. 2024. ‘HyperTraPS-CT: Inference and Prediction for Accumulation Pathways with Flexible Data and Model Structures’. bioRxiv, 2024–03.

Angaroni, Fabrizio, Kevin Chen, Chiara Damiani, Giulio Caravagna, Alex Graudenzi, and Daniele Ramazzotti. 2022. ‘PMCE: Efficient Inference of Expressive Models of Cancer Evolution with High Prognostic Power’. Bioinformatics 38 (3): 754–62.

Beerenwinkel, Niko, Martin Däumer, Tobias Sing, Jörg Rahnenführer, Thomas Lengauer, Joachim Selbig, Daniel Hoffmann, and Rolf Kaiser. 2005. ‘Estimating HIV Evolutionary Pathways and the Genetic Barrier to Drug Resistance’. The Journal of Infectious Diseases 191 (11): 1953–60.

Beerenwinkel, Niko, Roland F. Schwarz, Moritz Gerstung, and Florian Markowetz. 2015. ‘Cancer Evolution: Mathematical Models and Computational Inference’. Systematic Biology 64 (1): e1–25.

Boyko, James D., and Jeremy M. Beaulieu. 2021. ‘Generalized Hidden Markov Models for Phylogenetic Comparative Datasets’. Methods in Ecology and Evolution 12 (3): 468–78. 10.1111/2041-210X.13534.

Boyko, James D, Brian C O’Meara, and Jeremy M Beaulieu. 2023. ‘A Novel Method for Jointly Modeling the Evolution of Discrete and Continuous Traits’. Evolution 77 (3): 836–51. 10.1093/evolut/qpad002.

Caravagna, Giulio, Ylenia Giarratano, Daniele Ramazzotti, Ian Tomlinson, Trevor A. Graham, Guido Sanguinetti, and Andrea Sottoriva. 2018. ‘Detecting Repeated Cancer Evolution from Multi-Region Tumor Sequencing Data’. Nature Methods 15 (9): 707–14. 10.1038/s41592-018-0108-x.

Casali, Nicola, Vladyslav Nikolayevskyy, Yanina Balabanova, Simon R. Harris, Olga Ignatyeva, Irina Kontsevaya, Jukka Corander, et al. 2014. ‘Evolution and Transmission of Drug-Resistant Tuberculosis in a Russian Population’. Nature Genetics 46 (3): 279–86. 10.1038/ng.2878.

Csardi, Gabor, and Tamas Nepusz. 2006. ‘The Igraph Software Package for Complex Network Research’. InterJournal, Complex Systems 1695 (5): 1–9.

Diaz-Uriarte, Ramon, and Pablo Herrera-Nieto. 2022. ‘EvAM-Tools: Tools for Evolutionary Accumulation and Cancer Progression Models’. Bioinformatics 38 (24): 5457–59.

Diaz-Uriarte, Ramon, and Iain G. Johnston. 2024. ‘A Picture Guide to Cancer Progression and Monotonic Accumulation Models: Evolutionary Assumptions, Plausible Interpretations, and Alternative Uses’. arXiv. 10.48550/arXiv.2312.06824.

Felsenstein, Joseph. 1973. ‘Maximum Likelihood and Minimum-Steps Methods for Estimating Evolutionary Trees from Data on Discrete Characters’. Systematic Biology 22 (3): 240–49. 10.1093/sysbio/22.3.240.

Greenbury, Sam F., Mauricio Barahona, and Iain G. Johnston. 2020. ‘HyperTraPS: Inferring Probabilistic Patterns of Trait Acquisition in Evolutionary and Disease Progression Pathways’. Cell Systems 10 (1): 39–51.

Holt, Kathryn E., Heiman Wertheim, Ruth N. Zadoks, Stephen Baker, Chris A. Whitehouse, David Dance, Adam Jenney, et al. 2015. ‘Genomic Analysis of Diversity, Population Structure, Virulence, and Antimicrobial Resistance in Klebsiella Pneumoniae, an Urgent Threat to Public Health’. Proceedings of the National Academy of Sciences 112 (27): E3574–81. 10.1073/pnas.1501049112.

Johnston, Iain G., Till Hoffmann, Sam F. Greenbury, Ornella Cominetti, Muminatou Jallow, Dominic Kwiatkowski, Mauricio Barahona, Nick S. Jones, and Climent Casals-Pascual. 2019. ‘Precision Identification of High-Risk Phenotypes and Progression Pathways in Severe Malaria without Requiring Longitudinal Data’. NPJ Digital Medicine 2 (1): 63.

Johnston, Iain G., and Ellen C. Røyrvik. 2020. ‘Data-Driven Inference Reveals Distinct and Conserved Dynamic Pathways of Tool Use Emergence across Animal Taxa’. Iscience 23 (6).

Johnston, Iain G., and Ben P. Williams. 2016. ‘Evolutionary Inference across Eukaryotes Identifies Specific Pressures Favoring Mitochondrial Gene Retention’. Cell Systems 2 (2): 101–11. 10.1016/j.cels.2016.01.013.

Kassambara, Alboukadel. 2020. ‘Ggpubr:”Ggplot2” Based Publication Ready Plots’. R Package Version 0.4. 0 438.

Knutsen, Turid, Vasuki Gobu, Rodger Knaus, Hesed Padilla-Nash, Meena Augustus, Robert L. Strausberg, Ilan R. Kirsch, Karl Sirotkin, and Thomas Ried. 2005. ‘The Interactive Online SKY/M-FISH & CGH Database and the Entrez Cancer Chromosomes Search Database: Linkage of Chromosomal Aberrations with the Genome Sequence’. Genes, Chromosomes and Cancer 44 (1): 52–64. 10.1002/gcc.20224.

Lewis, Paul O. 2001. ‘A Likelihood Approach to Estimating Phylogeny from Discrete Morphological Character Data’. Systematic Biology 50 (6): 913–25.

Louca, Stilianos, and Michael Doebeli. 2018. ‘Efficient Comparative Phylogenetics on Large Trees’. Bioinformatics 34 (6): 1053–55. 10.1093/bioinformatics/btx701.

Louca, Stilianos, and Matthew W Pennell. 2020. ‘A General and Efficient Algorithm for the Likelihood of Diversification and Discrete-Trait Evolutionary Models’. Systematic Biology 69 (3): 545–56. 10.1093/sysbio/syz055.

Luo, Xiang Ge, Jack Kuipers, and Niko Beerenwinkel. 2023. ‘Joint Inference of Exclusivity Patterns and Recurrent Trajectories from Tumor Mutation Trees’. Nature Communications 14 (1): 3676. 10.1038/s41467-023-39400-w.

Moen, Marcus T., and Iain G. Johnston. 2023. ‘HyperHMM: Efficient Inference of Evolutionary and Progressive Dynamics on Hypercubic Transition Graphs’. Bioinformatics 39 (1): btac803.

Montazeri, Hesam, Jack Kuipers, Roger Kouyos, Jürg Böni, Sabine Yerly, Thomas Klimkait, Vincent Aubert, Huldrych F. Günthard, Niko Beerenwinkel, and Swiss HIV Cohort Study. 2016. ‘Large-Scale Inference of Conjunctive Bayesian Networks’. Bioinformatics 32 (17): i727–35.

Pagel, Mark. 1994. ‘Detecting Correlated Evolution on Phylogenies: A General Method for the Comparative Analysis of Discrete Characters’. Proceedings of the Royal Society of London. Series B: Biological Sciences 255 (1342): 37– 45.

Paradis, Emmanuel, and Klaus Schliep. 2019. ‘Ape 5.0: An Environment for Modern Phylogenetics and Evolutionary Analyses in R’. Bioinformatics 35 (3): 526–28. 10.1093/bioinformatics/bty633.

Peach, Robert L., Sam F. Greenbury, Iain G. Johnston, Sophia N. Yaliraki, David J. Lefevre, and Mauricio Barahona. 2021. ‘Understanding Learner Behaviour in Online Courses with Bayesian Modelling and Time Series Characterisation’. Scientific Reports 11 (1): 2823.

Pedersen, Thomas Lin. 2020. ‘Ggraph: An Implementation of Grammar of Graphics for Graphs and Networks’. R Package Version 2 (3): 1.

R Core Team, A., and R. Core Team. 2022. ‘R: A Language and Environment for Statistical Computing. R Foundation for Statistical Computing, Vienna, Austria. 2012’.

Revell, Liam J. 2012. ‘Phytools: An R Package for Phylogenetic Comparative Biology (and Other Things)’. Methods in Ecology and Evolution, no. 2, 217–23.

Schill, Rudolf, Maren Klever, Kevin Rupp, Y. Linda Hu, Andreas Lösch, Peter Georg, Simon Pfahler, et al. 2024. ‘Reconstructing Disease Histories in Huge Discrete State Spaces’. KI - Künstliche Intelligenz, January. 10.1007/s13218-023-00822-9.

Schill, Rudolf, Stefan Solbrig, Tilo Wettig, and Rainer Spang. 2020. ‘Modelling Cancer Progression Using Mutual Hazard Networks’. Bioinformatics 36 (1): 241–49.

Schliep, Klaus Peter. 2011. ‘Phangorn: Phylogenetic Analysis in R’. Bioinformatics 27 (4): 592–93. 10.1093/bioinformatics/btq706.

Schwartz, Russell, and Alejandro A. Schäffer. 2017. ‘The Evolution of Tumour Phylogenetics: Principles and Practice’. Nature Reviews Genetics 18 (4): 213–29. 10.1038/nrg.2016.170.

Wickham, Hadley. 2016. Ggplot2: Elegant Graphics for Data Analysis. Springer-Verlag. 10.1007/978-0-387-98141-3.

Yu, Guangchuang, David K. Smith, Huachen Zhu, Yi Guan, and Tommy Tsan-Yuk Lam. 2017. ‘Ggtree: An r Package for Visualization and Annotation of Phylogenetic Trees with Their Covariates and Other Associated Data’. Methods in Ecology and Evolution 8 (1): 28–36. <10.1111/2041-210X.12628>.

